# The Temporal Profile of Dual-task Interference in the Human Brain

**DOI:** 10.1101/2023.11.06.565914

**Authors:** Seyed-Reza Hashemirad, Maryam Vaziri-Pashkam, Mojtaba Abbaszadeh

**Affiliations:** School of Cognitive Sciences, Institute for Research in Fundamental Sciences (IPM), Tehran, Iran; Department of Psychological and Brain Sciences, University of Delaware, Newark, DE, USA; Department of Neurosciences, Université de Montréal, Montréal, QC, Canada

**Keywords:** Dual-task interference, Electroencephalogram (EEG), Multivariate analysis (MVPA), Drift diffusion modeling (DDM)

## Abstract

Due to the brain’s limited cognitive capacity, simultaneous execution of multiple tasks can lead to performance impairments, mainly when the tasks occur closely in time. This limitation is known as dual-task interference. We aimed to investigate the time course of this phenomenon in the brain, utilizing a combination of EEG, multivariate pattern analysis (MVPA), and drift-diffusion modeling (DDM). Here, participants first performed a tone discrimination task, followed by a lane-change task with either short or long onset time differences (Stimulus Onset Asynchrony, SOA), in a simulated driving environment. As expected, the dual-task interference increased the second task’s (lane-change) reaction time. The DDM analysis indicated that this increase was attributable to changes in both the decision time and the post-decision time. Our MVPA findings revealed a decrease in decoding accuracy for the lane-change task in short SOA compared to both long SOA and single-task conditions throughout the trial, highlighting the presence of interference. Moreover, the temporal generalization analysis identified a significant interference effect in short SOA compared to long SOA and single-task conditions after ∼250 ms relative to stimulus onset. Additionally, the conditional generalization analysis showed a delayed response after ∼450 ms. Searchlight analysis illustrated the progression of this information reduction, starting in occipital, parietal, and parieto-occipital leads responsible for perceptual and central processing and then transferring to the frontal leads for mapping decisions onto motor actions. Consistent with the hybrid dual-task interference theory, our results suggest that the processing of the two tasks occurs in a partial parallel manner for the first few hundred milliseconds and primarily in the perceptual and decision-processing stages. Subsequently, another competition arises between the two tasks to route information to motor areas for execution, resulting in the second task’s serial processing and delay or lengthening.

## Introduction

In today’s fast-paced world, multitasking has become increasingly common. Given the limitations of our cognitive resources, involving two or more tasks simultaneously can decrease performance. This effect is called dual-task interference. A prevalent form of this effect is driving while engaged in secondary activities. With the widespread adoption of smartphones and in-vehicle technologies, drivers often find themselves managing several tasks while operating a vehicle, which raises significant concerns about road safety due to its detrimental impact on driving performance. Extensive research has highlighted the adverse effects of dual-task on driving abilities (Abbas-Zadeh et al., 2021; Levy et al., 2006; Strayer & Johnston, 2001; Welford, 1952). A critical aspect to examine is the temporal dynamics of dual-task interference while driving. Understanding when interference occurs and its duration is essential for addressing safety concerns. To explore these dynamics, we conducted a study utilizing Electroencephalogram (EEG), Multivariate Pattern Analysis (MVPA), and Drift-Diffusion Modeling (DDM) within a controlled simulated driving environment.

Previous EEG studies on dual-task interference have predominantly relied on univariate methods to explore the neural correlations of dual-task interference (Dell’Acqua et al., 2005; Matthews et al., 2006; Pratt et al., 2011; Sigman & Dehaene, 2008; Töllner et al., 2012). These methods have focused on analyzing either the peak or the time course of Event-Related Potentials (ERPs). However, these findings do not clarify whether the observed brain activity changes result from general factors like task difficulty, attentional variations, and mental effort or if they are specifically due to interference in processing the tasks. A practical method to address this constraint is to explore the information over time for each task individually across the entire brain using multivariate pattern analysis (MVPA) during dual-task performance (King & Dehaene, 2014; Marti et al., 2015; Rajaei et al., 2019). By adopting MVPA, researchers gain the ability to explore the temporal progression of information processing in the brain while dual-task interference occurs. For instance, this method enables decoding specific features related to a driving task (e.g., detecting the direction of a turn or lane change) based on the pattern of responses across brain sensors. Changes in decoding accuracy during dual-task interference can then be measured to assess alterations in the information content at different time points.

Current theories of dual-task interference typically acknowledge three stages of processing for each task: perceptual, central, and motor stages (Hibberd et al., 2013; Pashler & Johnston, 1989; Sternberg, 1969). However, there is an ongoing debate about how dual-task interference affects these processing stages. In the literature, two main theories attempt to explain the underlying mechanism of dual-task interference: the bottleneck theory and the central capacity-sharing theory. The bottleneck theory suggests that when two tasks occur closely together, the central processing stage of the second task is delayed until the central stage of the first task is completed. However, the perceptual and motor stages can occur in parallel (McCann & Johnston, 1992; Pashler, 1994; Pashler & Johnston, 1989; Ruthruff et al., 2001; Sigman & Dehaene, 2005, 2006; Tombu et al., 2011). On the other hand, the central capacity-sharing theory proposes that the central stages of both tasks can be processed in parallel, with the limited cognitive resources shared between the tasks in a graded manner. This theory also rules out interference during the perceptual and motor stages (Duncan, 1980; Huestegge & Koch, 2010; Tombu & Jolicœur, 2002; Tombu & Jolicœur, 2003). The core disagreement between these theories revolves around the central or decision-related stage. The serial bottleneck theory posits strict sequential processing of tasks. In contrast, the capacity-sharing theory suggests that the central stage can handle multiple tasks concurrently but with the challenge of allocating limited resources, potentially leading to a decline in performance or slowed reaction times.

Using drift-diffusion modeling (DDM), a framework for modeling the distinct processing stages involved in two-choice tasks (Gold & Shadlen, 2007; Ratcliff et al., 2016; Shadlen & Newsome, 2001), recent research has indicated that the dynamics of dual-task interference cannot be fully explained by either of the bottleneck and central capacity-sharing theories alone (Abbas-Zadeh et al., 2021; Zylberberg et al., 2012). As a result, hybrid theories of dual-task processing have been put forward (Abbas-Zadeh et al., 2021; Zylberberg et al., 2012). According to the hybrid theory, the central stages of both tasks may be processed in a partial-parallel manner, but a bottleneck exists when routing the decision to motor structures. This means that not only are the central processing stages affected by dual-task demands, but the interference also extends into the motor stages. While these theories have mainly relied on behavioral data, integrating EEG and MVPA offers an opportunity to test them using neural data, providing a more comprehensive understanding of dual-task processing and interference.

Marti et al. (2015) applied the MVPA method of time decomposition to MEG data. They discovered a more complicated dynamic for the interference. They showed that brain processes could operate in parallel for the first ∼450 milliseconds, but beyond that, processing of the first task is shortened while processing of the second task is either lengthened or postponed. Neither the central capacity-sharing nor the bottleneck theories can fully explain these findings, but the hybrid theory might be suited to explain these results. Notably, Marti et al. (2015) study trained a decoder to distinguish between the non-interfering dual-task condition and the distractor condition. The key distinguishing factor between these two conditions was the presence or absence of motor execution. In other words, their decoder primarily discriminated based on the motor execution, diminishing the impact of task-specific information over time. To address this limitation, our approach involves training a decoder specifically to discriminate between the two conditions of each task in both single- and dual-task conditions. This approach allows us to investigate how dual-task interference influences the information processing of each task individually across time.

To date, most experimental designs exploring dual-task interference have predominantly focused on simple tasks like visual discrimination (e.g., distinguishing objects, colors, or orientations) or tone discrimination tasks (e.g., identifying high and low pitches). However, the gap between these artificial tasks and real-world scenarios, such as driving, is substantial. In real-life situations, individuals frequently encounter more intricate and sequential motor movements that involve multiple steps, which contrasts with the simplicity of artificial tasks typically limited to a single step. Additionally, the priority assigned to different tasks differs significantly. In artificial experiments, neither task inherently outweighs the other in importance. Conversely, in real-world driving, driving holds a significantly higher priority, thereby diminishing the significance of the secondary task, even if presented before the driving stimulus.

Moreover, real-world environments involve continuous tasks, exposing individuals to many distractions. In contrast, most artificial tasks consist of discrete and isolated stimuli. To approach the complexities of real-life dual-task situations, we developed a dual-task paradigm within a simulated driving environment (Abbas-Zadeh et al., 2021; Abbaszadeh et al., 2023). By considering these critical factors, we aimed to bridge the gap between experimental and real-world conditions, thereby enhancing the practicality and relevance of our research.

Taken together, here, we used a dual-task paradigm in which a tone discrimination task was followed by a lane-change task either after a short gap (100ms later, short Stimulus Onset Asynchrony, SOA) or after a long gap (600ms later, long SOA). Using MVPA and EEG, we investigated the temporal dynamic of information about the driving lane-change direction in the single- and dual-task conditions. We fitted a drift-diffusion model on the behavioral data to estimate the duration of the central or decision stage of the lane-change task and compare it to the subsequent MVPA results. Additionally, we used searchlight analysis to localize multivariate effects, determine the time course of each electrode classification accuracy, and examine which electrodes contributed to the classification accuracy at each time step. Together, the results of MVPA and DDM and their relationship can provide helpful information about the time course of dual-task interference in the human brain and provide evidence to test the validity of the models of dual-task interference.

### Results Behavioral results

The mean reaction time and accuracy were compared between the short, the long SOA, and single-task conditions in each task (Figure 1B). Reaction times for the lane-change task were 782 ± 71, 541 ± 84, and 605 ± 53 (mean ± std, here and after) for short SOA, long SOA, and single lane-change conditions, respectively. As was predicted by the psychological refractory period, there was a significant effect of condition on the reaction time of the lane-change task (F (2,36) = 168.4, p < 0.0001). Subsequent post-hoc analysis using Tukey HSD for multiple comparison correction revealed that the short SOA had a significantly longer reaction time in comparison to both the long SOA (p < 0.0001) and the single lane-change (p < 0.0001) conditions, indicating strong dual-task interference. Furthermore, a significantly shorter reaction time was observed in the long SOA condition compared to the single lane-change condition (p < 0.001). This difference in reaction time may be attributed to the priming effect of the tone task in the long SOA condition, which was not present in the single lane-change condition.

**Figure 1.**
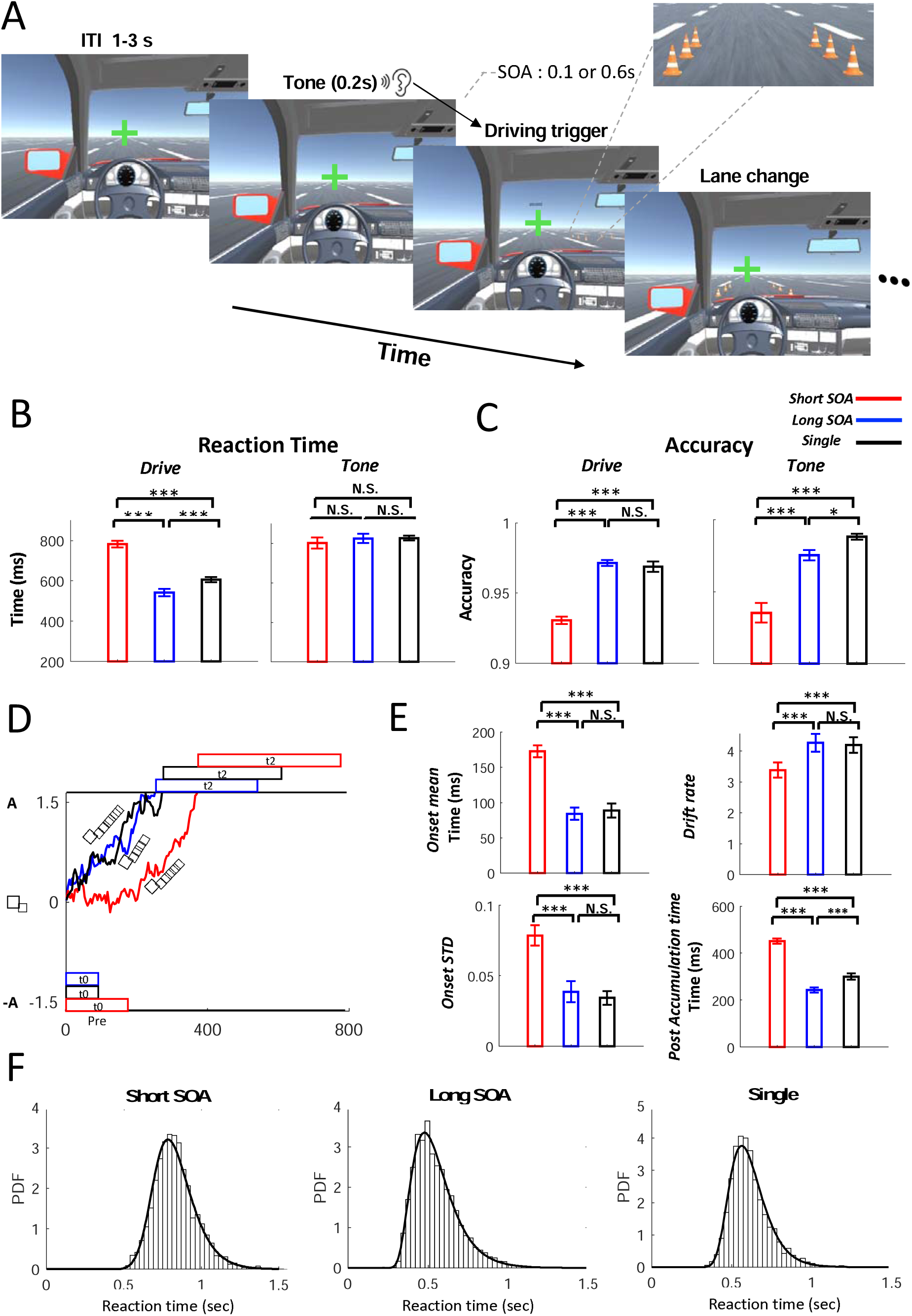
Paradigm, behavioral, and DDM results. A) Dual-task paradigm: The sequence of events for one sample trial. First, the tone task was presented and lasted for 200 ms, then the cones were presented with 100 or 600 ms delay (either long or short SOA) after the tone onset. The inter-trial interval (ITI) varied between 1 to 3 sec. Participants were instructed to respond to the tone task first and then to perform the lane-change as fast as possible. B) Reaction times (RT) and C) Accuracies for both tasks. The left and right panels show the results for lane-change and tone, respectively, for the short SOA (red), long SOA (blue), and single task (black) conditions. The error bars represent the standard error of mean (SEM, * p < 0.05, ** p < 0.01, *** p < 0.001). D) Schematic of drift-diffusion modeling based on estimated parameters by the model. A and X_O_ are the boundary and the starting point, and ^µ^ is the drift rate. t_0_ is the estimated onset of cumulative Gaussian that discounts early samples, and t_2_ is the post-accumulation time. Subjects accumulate evidence continuously until one of the decision boundaries is reached. Note, the reduction in drift rate coupled with increased post-accumulation time in short SOA that favors the hybrid model. E) Estimated parameters of the DDM for the lane-change task. Drift rate, Onset, STD, and post-accumulation times are presented for the long SOA (blue), short SOA (red), and single task (black) conditions. E) The RT distribution of the lane-change task for the three conditions for all the participants (histogram) and the model (black lines). The Bin width is 40 ms. The fraction of variance explained by the model fit (R^2^) was > 0.9 for all three conditions.

The lane-change task accuracy was 93 ± 1 %, 97 ± 0.8 %, and 96 ± 1% for short SOA, long SOA, and single lane-change conditions, respectively (Figure 1C). Analysis of lane-change task accuracy revealed a significant difference among conditions (F (2,36) = 142.31, p < 0.0001). Participants also had significantly higher driving accuracy in the long SOA (p<0.0001) and single drive (p < 0.0001) conditions in comparison to the short SOA condition; however, there was no significant difference between the accuracies of the single lane-change and the long SOA conditions (p = 0.51). The average lane-change accuracy was more than 90% in all three conditions.

Tone task reaction times were 800 ± 123, 825 ± 99, and 827 ± 103 for the short SOA, the long SOA, and the single tone conditions, respectively (Figure 1B). There was no significant difference between the reaction time of the tone task conditions (F (2,36) = 1.23, p = 0.3) because this task was always presented first. On the other hand, there was a significant difference between conditions for the tone accuracy (F (2,36) = 50.07, p < 0.001). The tone task’s accuracy was 93 ± 2 %, 97 ± 1 %, and 98 ± 0.9 % for the short SOA, the long SOA, and the single tone conditions, respectively (Figure 1C). The short SOA’s tone accuracy was significantly lower than the long SOA (p < 0.0001) and the single-tone condition (p < 0.0001). The long SOA also had significantly lower accuracy than the single-tone condition (p = 0.02).

These findings, consistent with our previous studies (Abbas-Zadeh et al., 2021; Abbaszadeh et al., 2023), are in agreement with the previous reports of dual-task interference in that performing two tasks in close succession significantly impairs the reaction time of the second task and the accuracy of both tasks, while the reaction time of the first task remains unaffected. Furthermore, the significantly lower reaction time of the long SOA compared to the single lane-change reveals the priming effect of the tone task in which the tone works as a cue to draw attentional resources to the expected lane-change task.

### Drift Diffusion Model

To better understand the effect of dual-task interference on the time course of each stage of the processing of the lane-change task, a drift-diffusion model (DDM) was fitted to the behavioral RTs of the lane-change task. DDM assumes that participants accumulate information continuously until sufficient evidence is gathered in favor of one of the choices (one of two thresholds is hit, Figure 1D)

For each participant, outliers were removed first, and then for each condition, four parameters (including means and standard deviations for the onset, drift rate, and post-accumulation time; see Methods for details) were considered. The starting point, boundaries, and sigma were considered equal across conditions. Subsequently, RTs of all conditions were pooled together, and the fitting procedure was performed only once for each subject using the maximum likelihood method. Finally, one-way repeated measures ANOVA was used to infer significance. The estimated parameters and R-squared for each participant are summarized in supplementary tables 1-3. The model was fit for each participant separately, and the average R-squared was > 0.9 for all three conditions (Figure 1F).

The mean estimated parameters are shown in Figure 1E. The mean drift rate during the accumulation time was 3.38 ± 1, 4.25 ± 1, and 4.18 ± 1 for the short SOA, the long SOA, and the single task conditions, respectively. There was a significant effect of condition on drift rate (F (2,36) = 15.54, p < 0.0001). Subsequent Tukey’s HSD post-hoc analysis revealed that both the long SOA and the single lane-change condition had significantly higher drift rates compared to the short SOA (ps < 0.001), however, there was no significant difference between the long SOA and the single lane-change condition (p = 0.9). The mean onset time was 172 ± 37, 84 ± 37, and 88 ± 43 for the short SOA, the long SOA, and the single condition, respectively. There was also a significant effect of the condition on the onset time mean (F (2,36) = 38.7, p < 0.0001). Consistent with drift rate results, both the long SOA and the single task had a shorter onset time than the short SOA (ps < 0.0001), while there was no significant difference between the long SOA and the single lane-change (p = 0.9). This may be due to a delay in processing at the beginning of the task caused by the secondary task in the short SOA condition. Moreover, the same effect was evident in the model’s standard deviation (std) of cumulative Gaussian. There was a significant effect of task condition on standard deviation (F (2,36) = 18.8, p < 0.0001). Likewise, both the long SOA and the single lane-change conditions had lower stds than the short SOA condition (long: p = 0.001; single: p < 0.001), implying a quick rise in the onset of evidence accumulation in these conditions compared to the short SOA. There was also no significant difference between the long SOA and the single lane-change in std (p = 0.88). Since the effect of drift rate and cumulative Gaussian parameters (mean and std of onset time) on conditions were similar, we will refer to these collectively as the evidence accumulation time.

Finally, the model estimated post-accumulation time was 451 ± 45, 242 ± 47, and 300 ± 56 ms for the short SOA, the long SOA, and the single lane-change conditions, respectively. There was a significant effect of task condition on post-accumulation time (F (2,36) =128, p < 0.0001). In post-hoc analysis, once again both the long SOA and the single lane-change had significantly shorter post-accumulation time than the short SOA (ps < 0.0001); however contrary to the previous parameters, the long SOA also had significantly shorter post-accumulation time than the single lane-change (p < 0.001), which suggests that the potential priming effect of tone on the lane change response in the long SOA condition, mainly affected the post-accumulation time resulting in shorter reaction times.

Altogether, the DDM results suggest that not only was the evidence accumulation time affected by the dual-task interference, but the interference also resulted in longer post-accumulation time. Hence, these results are in agreement with hybrid models that both the decision and non-decision times are affected as the two tasks get closer in time.

### Multivariate pattern analysis

Multivariate pattern analysis (MVPA) was applied to EEG data to characterize the representational dynamics of the dual-task interference. Applying this method to each task separately allows for a detailed look at the temporal dynamics of the information for each task and how each task is influenced by dual-task interference.

For each participant and task, MVPA was employed using Support Vector Machine (SVM) classifiers to analyze the data. The objective was to investigate potential differences in how information about lane-change direction conditions (right vs. left) and tone frequency discrimination (high vs. low) was encoded across conditions over time. Then, we performed a generalization analysis across time or condition by training an SVM on a particular time bin and condition and then generalizing and testing it on other time bins or conditions, respectively.

### Decreased decoding accuracy for the lane-change task in the short SOA condition

Figure 2 illustrates the decoding accuracy of the lane-change task, both time-locked to the onset of the lane-change stimulus (Figure 2A) and time-locked to the lane-change reaction time (Figure 2B) for the three task conditions. We employed a right-sided Wilcoxon signed-rank test with FDR correction, setting q to 0.05 to identify time-points with significantly above-chance decoding accuracy. We compared the earliest time that the decoding accuracy was above chance to determine the decoding onset latency. As depicted in Figure 2A, the average decoding onset latency was 225 ± 143, 158 ± 40, and 128 ± 35 ms (mean ± std resampling) for short SOA, long SOA and single-task conditions, respectively. There was a significant effect of condition on onset between conditions (F (2,198) = 31.61, P < 0.0001). Subsequent post-hoc analysis showed that the single lane-change condition had significantly earlier onset latency than the short and the long SOA conditions (ps < 0.0001). The long SOA onset latency was also earlier than the short SOA (p < 0.0001). Decoding accuracy was then followed by a rapid increase, reaching its first maximum peak accuracy at 264 ± 78 ms, 229 ± 56 ms, and 242 ± 64 ms for the short SOA, the long SOA, and the single lane-change conditions, respectively. There was no significant effect of condition on the difference between the first peak latency of either of the conditions, highlighting no delay (F (2,34) = 3.1, P = 0.07, Greenhouse-Geisser corrected).

**Figure 2.**
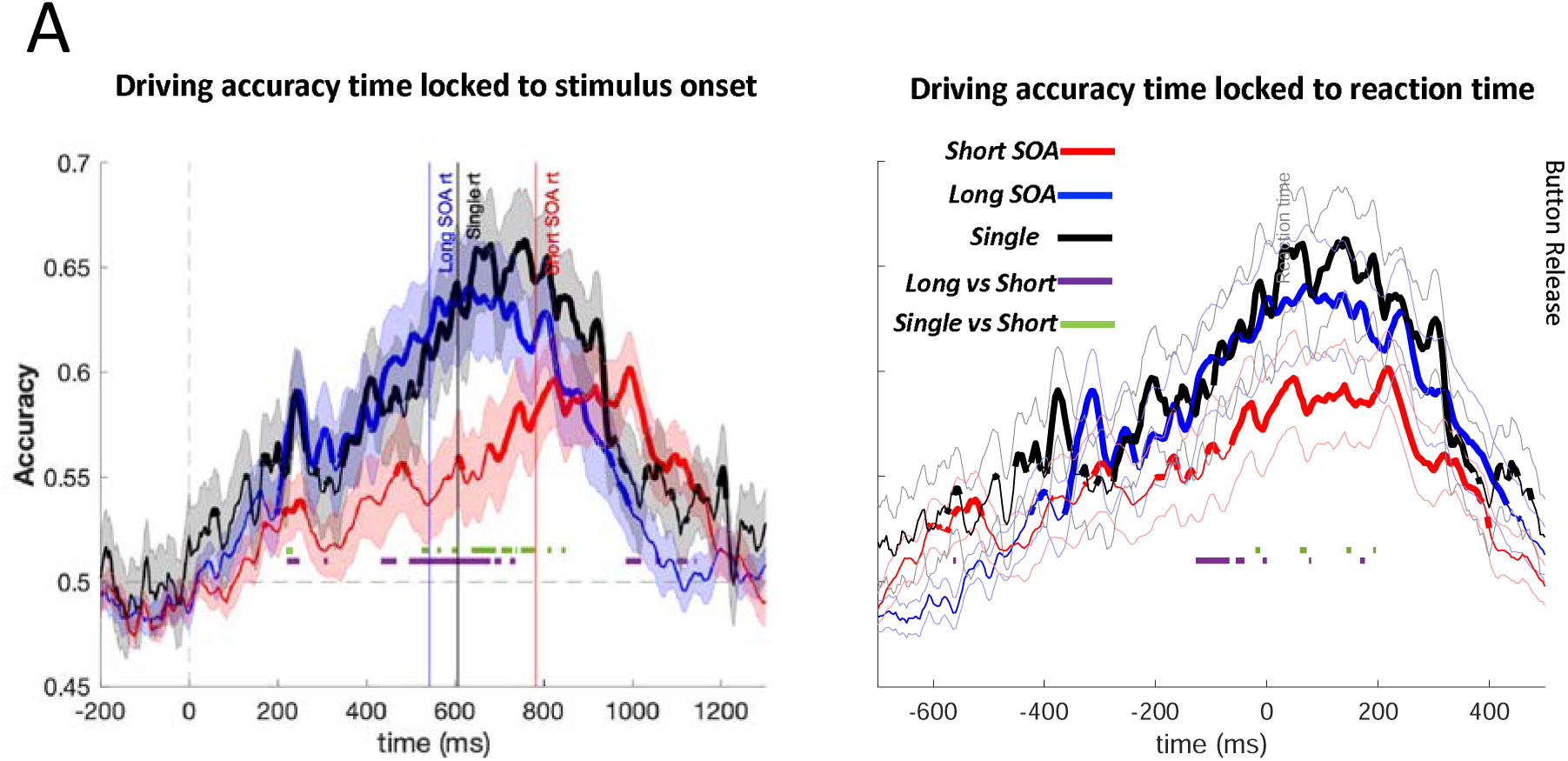
Temporal dynamics of dual task interference in lane-change task. A) Multivariate pattern analysis of EEG data. Time courses of decoding accuracies for the long, the short SOA, and the single lane-change condition averaged across 18 subjects time-locked to the stimulus onset (left) and the reaction time (right). Dashed vertical lines show lane-change onset. The chance level decoding accuracy is shown by the horizontal dashed line at 0.5. Three vertical lines are the corresponding reaction times of long, short SOA, and single drive conditions, respectively. Thicker lines indicate a decoding accuracy significantly above chance (right-sided signed-rank test, FDR corrected across time, q < 0.05), and shaded error bars represent the standard error of the mean (SEM). Ticker horizontal lines represent significant differences between long and short SOA (purple) and single drive and short SOA (green). There was no significant difference between the long SOA and the single drive decoding accuracy throughout the epoch. The legends indicate the corresponding reaction times and accuracies of each condition. For display purposes, data was smoothed using a moving average with five sample points.

Subsequently, a pairwise two-sided Wilcoxon signed-rank test was used to assess differences in decoding accuracy of the lane-change task conditions. The first period of difference in accuracies of the long SOA versus the short SOA started from 228 ms after stimulus onset, and it remained significant until 244 ms (Zs > 2.5, ps < 0.05). This was followed by a brief period from 432 ms to 468 ms (Zs > 2.54, ps < 0.042) and again around long SOA reaction time from 496 ms to 704 ms (Zs > 2.53, ps < 0.043). The next significant decoding accuracy periods were related to the movement time of the lane-change in the short SOA condition from ∼ 984 ms to 1128 (Zs > 2.53, ps < 0.043). Furthermore, there was no significant difference in the accuracy of the long SOA vs the single lane-change condition. The short SOA vs the single condition comprised of brief periods of significance: First brief periods from 48 ms to 60 ms and 236 ms to 244 ms (Zs > 2.5, ps < 0.05), followed by segregated periods from 524 ms to 784 ms before single drive movement time and finally from 836 ms to 864 ms associated to single drive movement time (Zs > 2.5, ps < 0.05).

As shown in Figure 2B, there was also a reduction in short SOA’s decoding accuracy compared to long SOA from ∼128 to 40 ms (Zs > 2.76, ps < 0.048) prior to the movement time and a very brief period from ∼20 to 10 ms (Zs > 3.2, ps < 0.035) compared to the single condition in reaction time-locked decoding accuracy. This reduction in decoding accuracy right before the movement once again suggests an interference during motor processing and execution stages.

On the other hand, there was no significant above-chance decoding accuracy in either of the long SOA, short SOA, and single conditions or their differences for the tone task (Figure S1). One reason for this low accuracy might be the relatively minor frequency difference between the two tone conditions compared to the pronounced left/right shift required for the lane change task.

Taken together, the significant reduction in the decoding accuracy in the short compared to the long SOA across the trial (and even a brief period in the short SOA vs the single lane-change) contradicts the pure bottleneck theory. This theory hypothesizes that when two tasks occur in close succession, the central processing of the second task is delayed until after the central stage of the first task. As a result, the decoding accuracy of the second task in the short SOA should show a delay and then a rise to the same level as the long SOA condition as the two tasks are being processed in serial; hence, this theory cannot explain the observed pattern of drop in decoding accuracy for the lane-change task in short SOA condition. On the other hand, the capacity sharing model does not predict any delay in processing and cannot explain the later rise in decoding accuracy in short compared to long SOA at the beginning of the trial in our results.

### Different patterns of generalization in the short compared to the long SOA condition

To evaluate how dual-task interference may affect the temporal generalization of each condition, time-time decoding matrices were created for each task condition. To construct the time-time decoding matrices, an SVM classifier trained at one time was tested on its discrimination performance at all other times of the given condition. Temporal generalization provides helpful information about how different stages of brain processes evolve and interact across time.

The temporal generalization pattern changed across conditions (Fig. 3A). The overall time-time decoding matrix in the long SOA represented two distinct patterns of generalizations. First, a diagonal-shaped generalization is evident (King & Dehaene, 2014) from ∼0-250 ms after stimulus onset. This pattern shows that a classifier trained on each time point can generalize to the test from the same time point and a few time points around it, leading to a region of orange-red in the figure in the green background that lays on the diagonal of the matrix seeping to regions slightly around the diagonal. This pattern suggests that the perceptual and central processing stages happen in a partially overlapping cascade. This was followed by a square-shaped pattern (King & Dehaene, 2014) of generalization that started ∼300 ms before reaction time until ∼500 ms after it (∼250 ms to ∼1000 ms after stimulus onset). This pattern shows that a classifier trained on each time point can generalize to most time points in this temporal duration, creating a square-shaped red region within the green background. This pattern suggests a sustained processing pattern in this period, probably due to sustained motor-related activities. This sustained generalization is reasonable even after the first button press reaction time, as the participants should hold the key and maintain the car position between the cones.

**Figure 3.**
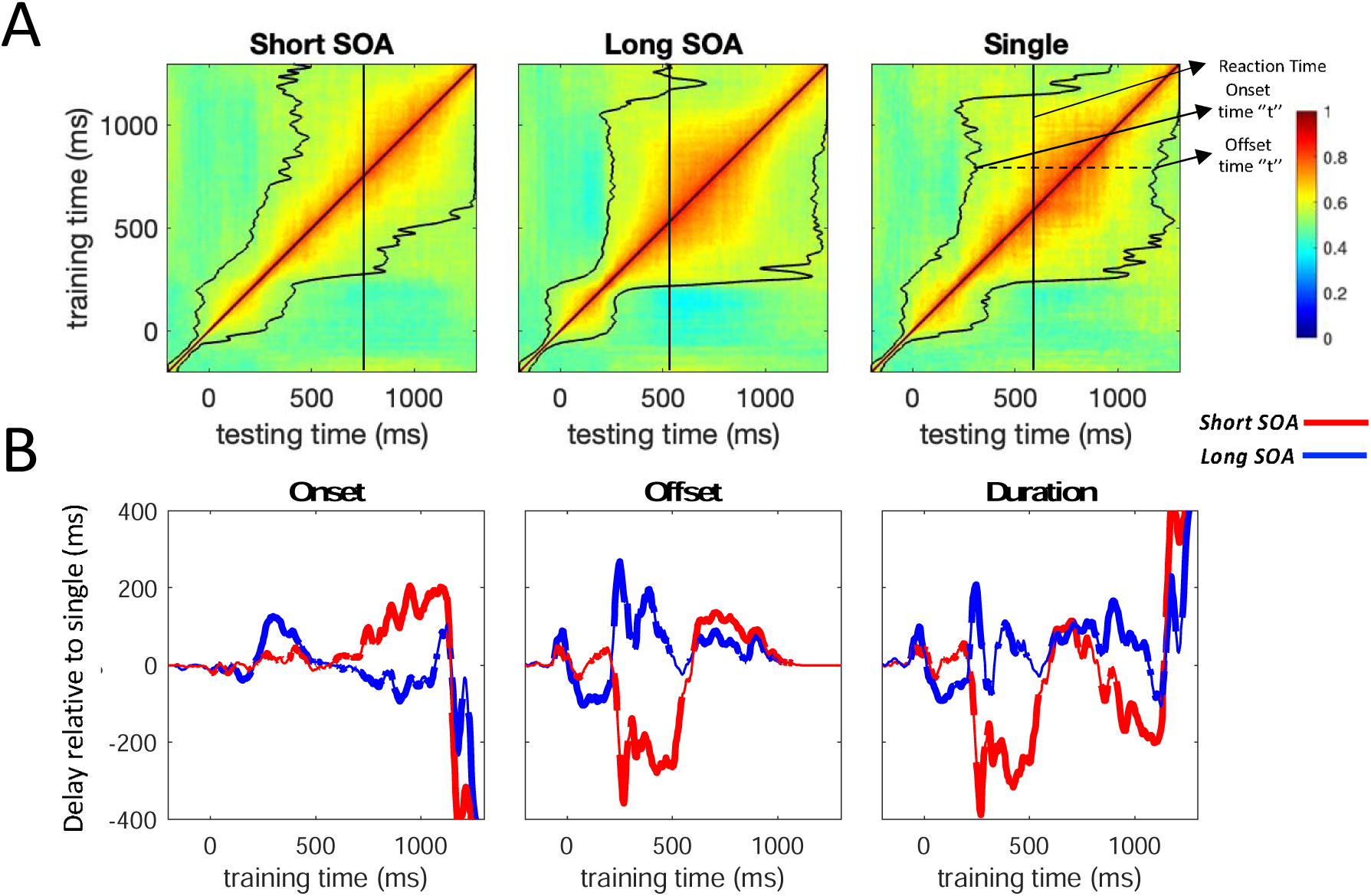
Temporal generalization patterns for the lane-change task. A) Time-time decoding matrices of the long SOA, the short SOA, and the single lane-change task. A classifier trained at any given time is also tested at all other time points, resulting in a training time x testing time temporal generalization matrix for each condition. The horizontal axis indicates testing times, and the vertical axis indicates training times. Color bars represent the percent of decoding AUC (chancel level = 50%). Within each time-time decoding matrix, onset and offset latencies are shown by solid black lines and smoothed by a moving average of five samples. **B)** Parameters of temporal generalization for the lane-change task. Onset latency was defined as the earliest time performance became significantly above chance (i.e., AUC=0.5) for at least five consecutive time bins (20 milliseconds). Offset latency was defined as the earliest time decoding performance was insignificant for 20 milliseconds. The standard deviation was computed by a resampling method. The duration was defined as the time between onset and offset latencies. Colored lines represent the mean difference between the single lane-change (horizontal dash line) and both the long (blue) and the short (red) SOAs. Positive and negative values indicate earlier and delayed onset and offset latencies and longer and shorter duration periods, respectively. Thicker lines represent a significant (two-sided signed-rank test, FDR corrected across time, q < 0.05) difference from the single lane-change condition. For display purposes, data was smoothed using a moving average with five sample points.

A similar pattern was also evident in the time-time decoding matrix of the single lane-change condition. However, a broader generalization pattern was noticeable compared to the long SOA, suggesting broader generalization across time in this condition in both central and motor stages.

For the short SOA, significantly above-chance decoding performance started right after stimulus onset with a diagonal generalization pattern. The distinct square-shape and diagonal-shape generalization patterns are not evident in short SOA, and the overall generalization is dampened. This suggests strong interference in this condition during the central and motor processing stages, particularly beyond ∼250ms.

The statistical comparison of the differences in temporal generalization between lane-change conditions was performed by calculating the onset, offset, and duration for generalization accuracy at each time point. At each training time, onset latency was defined as the earliest time where performance became significantly above chance (i.e., AUC > 0.5) for at least five consecutive time bins (20 milliseconds) across subjects. Likewise, offset latency for each training time was defined as the earliest time where performance became non-significant for at least five consecutive time bins, and finally, duration was the difference between offset and onset latencies (see methods). The single lane-change condition was considered the baseline condition, and the two other conditions were compared with this baseline for onset, offset, and duration parameters (Figure 3B).

Initially the classifiers trained in the first ∼200 ms from lane-change onset, the long SOA condition had earlier onset at ∼124 ± 41 to 196 ± 57 ms (Zs > 2.9, ps < 0.004) and earlier offset from ∼0 to 200 ms (Zs > 2.12, ps < 0.047) compared to the single lane-change condition, resulting in a shorter duration from 0 ± 42 to 200 ± 63 ms (Zs > 2.2, ps < 0.034). This indicates quicker processing of stimulus in long SOA that can later facilitate the earlier motor response in long SOA. Moreover, the short SOA condition had comparable onset, offset, and duration to the single lane-change condition with no delays in the initial 200 ms, suggesting parallel processing of tasks at the early stages.

Subsequently, in the long SOA, processing entered into the sustained generalization of motor-related activities that resulted in the delayed onset observed in classifiers trained at approximately 212 ± 8 to 480 ± 74 ms (Zs > 2.1, ps < 0.04), and a delay in offset latency from ∼200 ± 18 to 296 ± 177 ms and 344 ± 178 to 400 ± 204 ms compared to the single lane-change (Zs > 2.12, ps < 0.046). Additionally, classifiers trained from ∼200 ± 65 to 250 ± 27 and ∼ 400 to 500 ms exhibited significantly longer durations compared to the single condition (Zs > 2.01, ps < 0.05), suggesting that the sustained generalization (related to motor preparation/movement) occurred earlier for the long SOA and the processing happened longer in duration. The earlier start is consistent with the behavior and DDM results since long SOA had significantly shorter RT and also shorter post-accumulation time in the DDM model. Comparing short SOA and single condition, although there were no significant differences between them in lane-change onset latency in 200-600 ms, there was an earlier offset, from ∼220 ± 11 to 546 ± 56 ms after stimulus onset (Zs > 2.13, ps < 0.046), resulting in a shorter duration in this period. This generalization indicates competition between two tasks for decision-making and routing information to motor areas for execution.

Finally, the long SOA compared to the single condition showed an earlier onset in classifiers trained at ∼ 708 ± 71 to 1064 ± 72 ms (Zs > 2.07, ps < 0.049) and ∼1148 ± 22 to 1300 ± 32 ms (Zs > 2.1, ps < 0.04) and also delayed offset from ∼600 to 1000 ms compared to the single lane-change (Zs > 2.16, ps < 0.043), resulting in longer duration including classifiers trained at 600 ± 154 to 1100 ± 21 ms and 1200 ± 39 to 1300 ± 32 ms (Zs > 2.08, ps < 0.047), highlighting the more extensive sustained generalization related to motor movements. The short SOA condition, compared to the single condition, had a delayed onset from approximately 600 ± 58 ms to 1148 ± 308 ms (Zs > 2.15, ps < 0.04), which corresponds to the postponed motor execution. However, the short SOA showed an earlier onset in classifiers trained from 1152 ± 301 ms to 1300 ± 4 ms (Zs > 4.6, ps < 0.0001). This period was also accompanied by a significant delayed offset of ∼600 ± 105 to 1060 ± 74 ms (Zs > 2.1, ps < 0.044), beyond which time there was no difference in offset latency between the conditions. As such, there was a longer duration of generalization from 600 ± 154 to 720 ± 168 ms (Zs > 3.44, ps < 0.0001) followed by another shorter duration from 836 ± 143 to 1136 ± 222 (Zs > 2.17, ps < 0.035) ms around the time of motor execution.

These results emphasize the shorter duration and quicker processing of the long SOA before motor-related activities compared to both the short SOA and the single condition, leading to a faster transition to motor areas for execution. Notably, in the short SOA, the earlier offset and shorter duration from ∼250-600 ms compared to the single condition highlights the strong interference and competition for the two tasks for decision-making and routing information to motor areas that delay execution.

### Conditional generalization reveals a delay in the short SOA processes

To construct conditional generalization matrices, we employed classifiers trained in one condition and tested their performance in other conditions. This approach allowed us to investigate whether the observed temporal generalization differences between conditions were linked to a higher signal-to-noise ratio in the long SOA and single conditions or if they revealed distinct patterns of generalization and information processing speeds (Rajaei et al., 2019). The overall generalization displayed a narrow period of significance with a delayed onset and earlier offset (Figure 4A). This phenomenon might be attributed to varying time courses of processing in different conditions, which can affect decoding performance. To ensure a reliable comparison across conditions, we focused solely on the time of peak accuracy (peak latency) in the analysis (Figure 4B).

**Figure 4.**
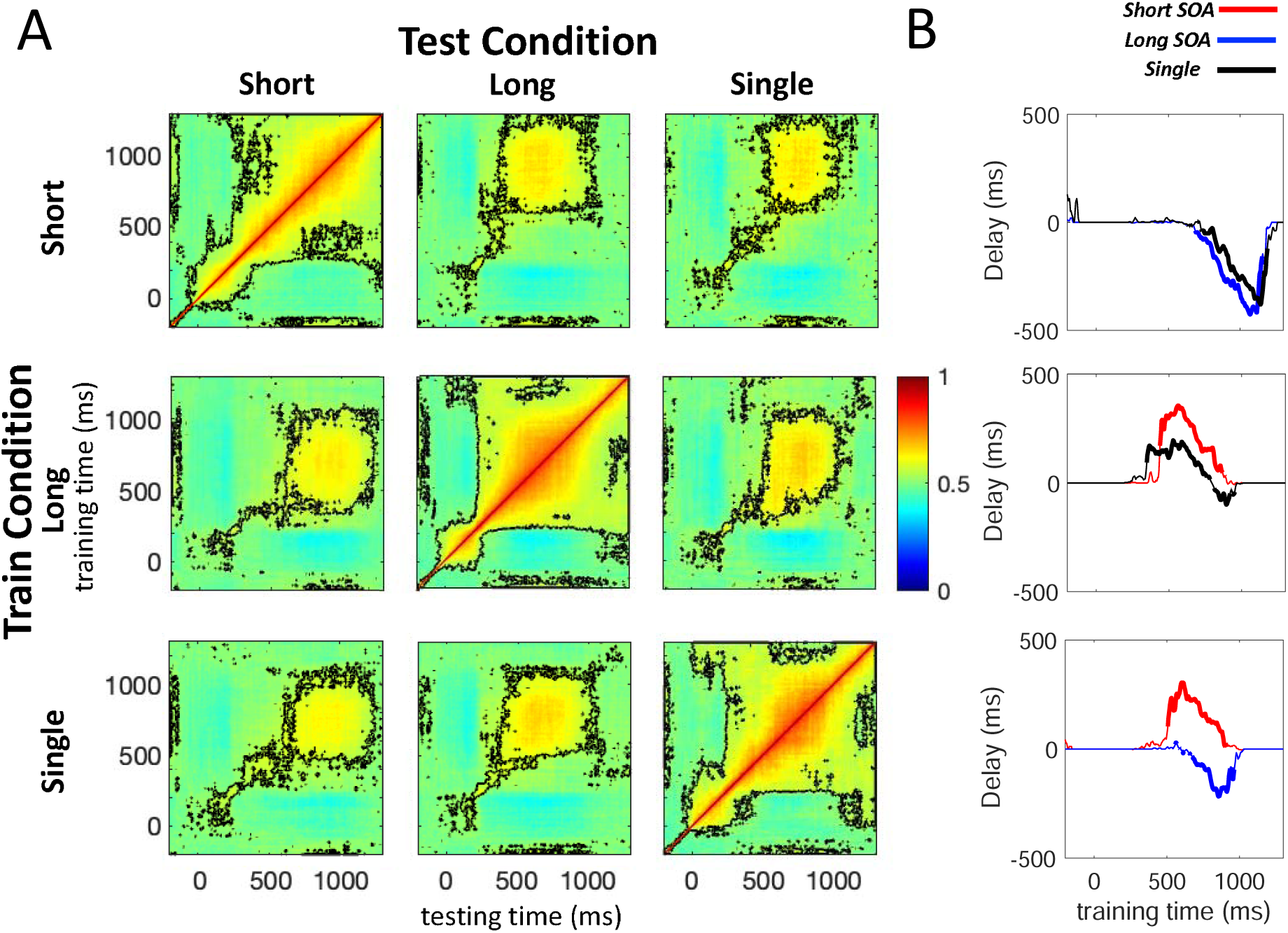
Conditional generalization of the lane-change task. A) Conditional generalization matrices. The classifiers were trained on one condition and then tested to discriminate right and left shifts in other conditions. Significantly above-chance decoding performance is shown by dashed contour lines (right-sided signed-rank test, FDR-corrected across time, q < 0.05). Each figure’s horizontal axis indicates testing times, and the vertical axis indicates training times. Each row represents the classifier training condition, and each column shows the testing condition. Color bars represent the percent of decoding AUC (chancel level = 50%); **B)** Peak latency parameter of conditional generalization matrix. Peak latency was considered as the time point where accuracy is maximum. In each row, the accuracy of the within-condition classification (e.g., long/long with training on long SOA and testing on long SOA) is represented as the horizontal dashed line. The resulting peak latency of the other two generalizations (e.g., long/single and long/short) are plotted relative to this horizontal line. Thicker lines represent a significant (two-sided signed-rank test, FDR corrected across time, q < 0.05) difference from the dashed line.

The short/long (train on short, test on long) and short/single (train on short, test on single) generalization matrices exhibited a narrow generalization pattern, displaying a delayed onset and an earlier offset throughout the time course of generalization (Figure 4A, top row). Compared to when training and test were both done on the short SOA condition (short/short), the peak latency occurred earlier in classifiers trained from 700 ± 41 to 1150 ± 233 ms for the short/long generalization (Zs > 3.5, ps < 0.001) and from 750 ± 67 to 1150 ± 224 ms for the short/single generalization (Zs > 3.1, ps < 0.004, Figure 4B, top panel). Consequently, the generalization was shifted towards the upper diagonal areas.

The time-time decoding matrix, trained in the long SOA condition and tested in the short SOA (long/short generalization), exhibited a delayed pattern of generalization compared to when training and test were both done on the long SOA condition (long/long Figure 4A, middle row). The long/short generalization matrix demonstrated narrow generalization, with a delayed onset and earlier offset in most cases. Notably, the peak latency was significantly delayed in classifiers trained from 444 ± 200 to 870 ± 68 ms (Zs > 3.6, ps < 0.0001), leading to a shifted generalization pattern below the diagonal (Figure 4B, middle panel). Similarly, the long/single generalization also displayed a delayed peak latency in classifiers from ∼348 ± 118 to 790 ± 45 ms but to a lesser degree (Zs > 3.6, ps < 0.0001) followed by an earlier latency from ∼800 to 900 ms (Zs > 3.6, ps < 0.0001, Figure 4B, middle).

Furthermore, the same generalization pattern was observed when training on the single condition (Figure 4A, bottom row). The single/short generalization matrix exhibited a narrow generalization pattern with a delayed peak latency in classifiers trained from ∼560 ± 130 to 890 ± 41 ms (Zs > 3.2, ps < 0.003) compared to the single/single generalization (Figure 4B, button panel). This delay in the peak latency is likely related to the motor execution process, causing the generalization matrix to shift below the diagonal. In the single/long generalization matrix, an earlier peak latency was observed in classifiers trained from 650 ± 24 to 960 ± 125 ms (Zs > 3.2, ps < 0.003), corresponding to the time of motor execution (Figure 4B, button panel). These conditional generalization results align with those of Marti et al. (2015), indicating a delay in the short SOA after ∼450 ms during motor processing.

### Searchlight analysis disclosed the topographical flow of information processing

A searchlight analysis was conducted to identify the most informative EEG electrodes related to the lane-change task. This analysis is a useful approach to localize multivariate effects; for each channel and its neighboring channels, decoding was performed using a sliding window of 50 ms as a feature matrix with steps of 4 ms. At the beginning of the trial, leads over the parietal and occipital regions were more involved in classification performance. As time elapsed, the information was gradually transferred to the frontal leads for response selection and motor execution. The decoding performance is shown in Figure 5 for the short SOA, the long SOA, and the single lane-change conditions in steps of 200 ms.

**Figure 5.**
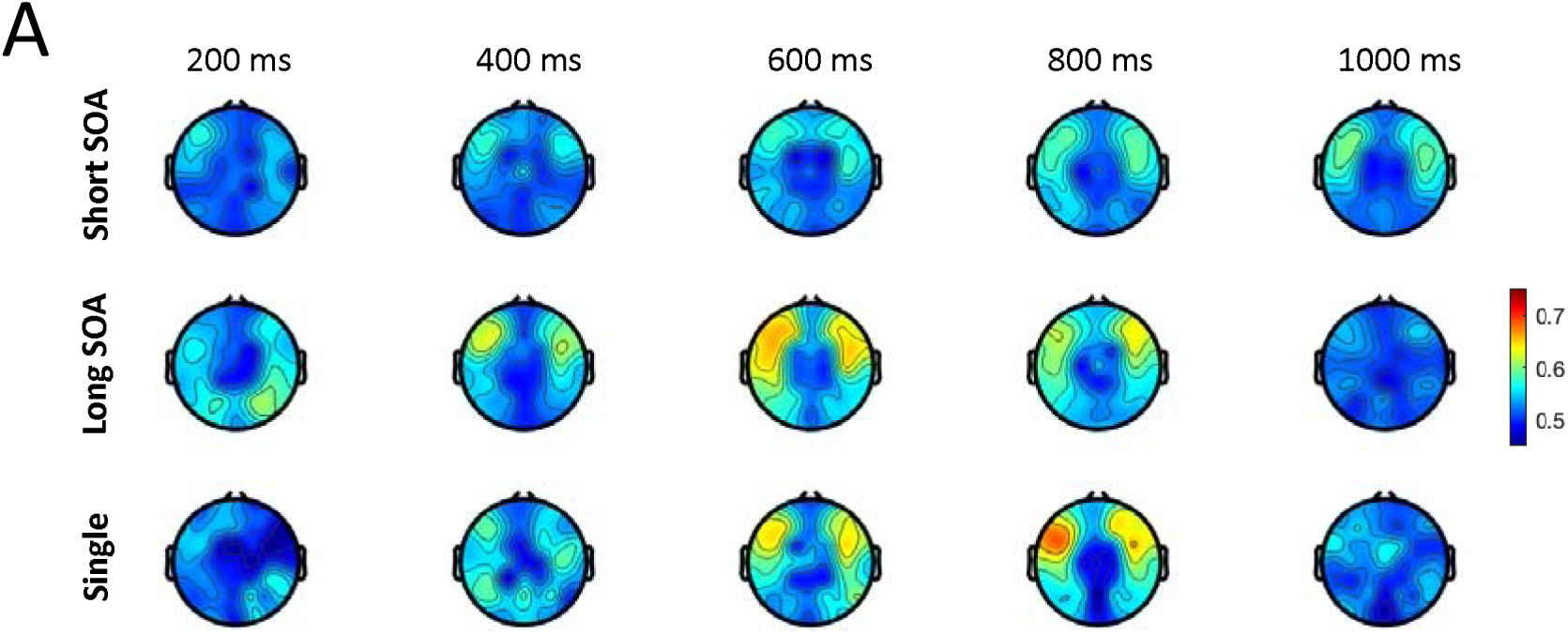
Searchlight analysis. Topographical plot of searchlight analysis for the short SOA (upper row), long SOA (middle row) and single lane-change condition (lower row) in steps of 200 ms. Searchlight analysis was done using a sliding window of 50 ms with 4 ms steps. In each window the decoding performance (AUC) of each electrode and its neighboring electrodes was calculated and the corresponding topographical map was plotted using EEGLAB ‘topoplot’ function.

In short SOA, performances were low throughout the trial. In 200 ms, only a few electrodes in the right centro-parietal (‘CP4’, ‘CP6’), frontal electrodes (‘F3’, ‘F4’, ‘F5’, ‘F7’, ‘AF7’), ‘FC5’, ‘TP8,10’ had significant decoding performance (Zs > 2.3, ps < 0.049). Notably, the maximum decoding was in left lateral frontal electrodes (Zs > 2.55, ps < 0.03), and decoding performance among parietal and occipital electrodes was compromised by interference compared to the long SOA. At 400 ms, only frontal electrodes (‘AF7’, ‘F3’, ‘F5’, ‘F6’, ‘F7’) had significant performance (Zs > 2.3, ps < 0.049).

In 600 ms, frontal electrodes’ performance remained significant (‘AF3’, ‘AF7’, ‘F1’, ‘F3’, ‘F5’). Moreover, fronto-central (‘FC4’), fronto-parietal (‘FP1’), central (‘C4’), centro-parietal (‘CP5’), parietal (‘P7’) and temporo-parietal (‘TP7’, ‘TP9’) had also significant performances (Zs > 2.3, ps < 0.049). At the time of reaction time around 800-1000 ms frontal electrodes had higher decoding performance (‘AF3’, ‘AF7’, ‘F1’, ‘F3’, ‘F4’, ‘F5’, ‘F6’, ‘F7’, ‘FC3’, ‘FC4’, ‘FC5’, ‘FC6’, ‘FT10’, ‘FT7’, (Zs > 2.65, ps < 0.03). Moreover, parietal, central, centro-parietal and parieto-occipital electrodes also had significant performance (‘C3’, ‘C4’, ‘C5’, ‘C6’, ‘CP3’, ‘CP4’, ‘CP6’, ‘P3’, ‘P7’, ‘PO3’, ‘PO7’, (Zs > 2.3, ps < 0.049) indicating a high decoding accuracy and extensive processing at the time of motor execution.

For the long SOA, at 200 ms after stimulus onset, parietal and occipital electrodes had higher decoding performance. All parietal and parieto-occipital electrodes’ performances were significant. Moreover, ‘O1’, lateral central and centro-parietal (‘C3’, C4’, ‘C5’, ‘C6’, ‘CP3’, ‘CP4’, ‘CP6’), lateral frontal and fronto-central (‘AF8’, ‘F4’, ‘F5’, ‘F6’, ‘FC3’, ‘FC4’, ‘FC5’, ‘FC6’), temporal and temporo-parietal electrodes (‘T7’, ‘T8’, ‘TP10’, ‘TP8’) had significantly above chance performance (Zs > 2.39, ps < 0.042) with maximum decoding in parietal (‘P4’, ‘P8’, ‘P6’), parieto-occipital (‘PO4’, ‘PO8’) and centro-parietal electrodes (‘CP4’, ‘CP6’, Zs > 2.7, ps < 0.025). After that in 400 ms, frontal electrodes had higher decoding performance indicating the preparation of information for execution. All frontal electrodes, lateral fronto-central (‘FC3’, ‘FC4’, ‘FC5’, ‘FC6’), fronto-temporal (‘FT10’, ‘FT7’, ‘FT8’), parietal (‘P6’, ‘P7’) and temporo-parietal (‘TP7’, ‘TP10) had significant performance (Zs > 2.3, ps < 0.049), with maximum decoding in frontal (‘F3’, ‘F5’, ‘F7’) and fronto-central (‘FC3’, ‘FC4’, ‘FC5’, ‘FC6’) electrodes (Zs > 3.04, ps < 0.016).

Likewise, in 600 and 800 ms, the same frontal, lateral fronto-central, fronto-parietal (‘FPz’, ‘FP1’) and fronto-temporal (‘FT10’, ‘FT7’, ‘FT8’) electrodes had the highest performance (Zs > 2.36, ps < 0.045). Moreover, lateral central, all parietal and parieto-occipital electrodes, temporal and temporo-parietal electrodes (‘T7’, ‘T8’, ‘TP10’, ‘TP7’, ‘TP9’) had significantly above chance performance (Zs > 2.36, ps < 0.045). However, the performance gradually decreased since the trial was nearly completed and in around 1000 ms few electrodes in lateral frontal (‘AF7’, ‘F4’, ‘F5’, ‘F6’, ‘F7’), centro-parietal (‘CP3’, ‘CP5’), ‘FP2’, ‘FC3’, ‘TP7’ and ‘TP9’ had significant performance (Zs > 2.35, ps < 0.046).

The single lane-change followed a similar long SOA decoding time-course but with reduced decoding performance. In 200 ms, only a few electrodes in the right lateral centro-parietal (‘CP4’, ‘CP6’) had significantly above chance decoding (Zs > 2.25, ps < 0.05). Notably, in 400 ms centro-parietal (‘CP3’, ‘CP5’), parietal (‘P3’, ‘P5’), temporo-parietal (‘TP10’, ‘TP7’, ‘TP9’), left lateral frontal (‘F3’, ‘F5’) and fronto-temporal (‘FT10’) had significant performance (Zs > 2.39, ps < 0.042), whereas in the long SOA, the information processing was most significant in frontal electrodes in time 400 ms. In 600 ms, frontal electrodes had higher performance. All frontal electrodes, lateral fronto-central (‘FC3’, ‘FC4’, ‘FC5’, ‘FC6’), fronto-temporal (‘FT10’), right lateral parietal (‘P6’, ‘P8’) and central (‘C1’, C4’) electrodes had significantly above chance decoding performance (Zs > 2.36, ps < 0.045). In 800 ms, the processing was more extensive as all frontal electrodes, lateral fronto-central (‘FC3’, ‘FC4’, ‘FC5’, ‘FC6’), fronto-temporal (‘FT10’, ‘FT7’, ‘FT8’), lateral central and centro-parietal (‘C4’, ‘C5’, ‘CP3’, ‘CP4’, ‘CP6’) had significant performance (Zs > 2.31, ps < 0.049) with maximum performance in left lateral frontal electrodes (Zs > 3.02, ps < 0.016). Once again, in 1000 ms, performance declined, and only a few electrodes (‘FC5’, ‘FCz’, and ‘FT7’) had significant performance (Zs > 2.43, ps < 0.039).

Taken together, the topographical time-course clearly suggests the pattern of information processing that begins in the occipital, parietal, and parieto-occipital leads responsible for visual response and decision-making and then transfers to the frontal electrodes for mapping the decision on the motor regions. The decoding performance was significantly compromised in the short SOA. Moreover, the strong interference in temporal generalization of short SOA (beyond ∼250 ms) occurred primarily in frontal and fronto-central electrodes. However, these speculations should be considered with caution as we did not perform source localization on our data.

## Discussion

This study aimed to investigate the time course of information processing during dual-task interference in the brain through a combination of EEG, multivariate pattern analysis (MVPA), and drift-diffusion model (DDM). We aimed to test different theories proposed to explain the dual-task paradigm, mainly capacity sharing, bottleneck, and hybrid models, and to find a model that could best account for the dynamics of dual-task interference. Our behavioral results, consistent with our previous studies (Abbas-Zadeh et al., 2021; Abbaszadeh et al., 2023), showed an increase in RT in the second task and a decrease in accuracy of both tone and lane-change task in the short SOA compared to the long SOA trials, confirming that the dual-task interference influenced the performance. The EEG results showed a decline in driving information during the short compared to long SOA and single conditions. Furthermore, the interference in the short SOA condition resulted in a different and dampened pattern of temporal generalization and a delay beyond ∼450ms in conditional generalizations compared to long and non-interfering SOA condition. Eventually, the searchlight analysis revealed a decline in the decoding accuracy of short SOA in electrodes spanning a wide range of brain regions.

The decoding accuracy of the lane-change task in the short SOA was significantly reduced compared to both the single and the long SOA conditions. These results are not consistent with pure bottleneck theory, which predicts the central processing resources can only be made available for one task at a time (McCann & Johnston, 1992; Ruthruff et al., 2001; Sigman & Dehaene, 2005, 2006; Tombu et al., 2011); on the contrary, our data showed that the processing of two tasks occurs concurrently but with reduced efficiency, indicating partial parallel processing. The same effect was also evident in the DDM results, as the model predicted a reduced rate of evidence accumulation. Moreover, the results revealed a delayed onset in both DDM and onset latency of MVPA in the short SOA, suggesting an interference emerging as soon as the second task begins, causing a delay in the onset of evidence accumulation. This also indicates that a pure capacity sharing model cannot account for the results. The neural and modeling results suggest that a hybrid model better accounts for the entirety of the results.

In the temporal generalization analysis, the generalization matrix showed a different pattern in the short SOA condition compared to both the single and the long SOA conditions. This difference was primarily related to the processing stages in classifiers trained beyond ∼250 ms (Figure 3,4A) stretching to ∼600 ms. The decoding accuracy in the main MVPA also significantly dropped around the same time. This period mainly points to the interference related to the central processing (decision-making) stage as well as the routing of information to motor areas for execution. In the searchlight analysis, the spatial interference pattern in this period is mainly observed in frontal and front-central electrodes. Furthermore, the DDM also revealed both a slow accumulation rate and a longer post-accumulation time in short SOA compared to long and single conditions, which is consistent with the pattern of interference in EEG being stretched all the way to the execution of motor movement. Taken together, these results once again contradicted the conventional hypothesis that only the central stages can be affected by interference (Pashler & Johnston, 1989) and were in favor of the hybrid theory (Abbas-Zadeh et al., 2021; Zylberberg et al., 2012) that a degree of interference was evident in accumulation and post-accumulation times. As mentioned before, Zylberberg et al. (2012) and Abbas-Zadeh et al. (2021) suggested a hybrid model for dual-task interference where the central stage of two tasks can be processed in parallel. However, interference also exists to route the decision to motor structures.

Our conditional generalization results revealed that when trained on the long condition and tested on the short SOA condition, the peak latency of decoding accuracy was delayed compared to when the training and testing were performed within the long SOA condition. This delay was observed in training times around 450 ms, stretching to training times around 870 ms(Figure 4B). These results are consistent with the previous report (Marti et al., 2015) and suggest that although the processing is affected before 450 ms (as discussed above), there is no delay in processing the second task.

However, beyond 450 ms, the interference causes a delay or lengthening of the processing of the second task. These results are also consistent with previous studies on event-related potentials (ERPs) looking at P300, which have demonstrated a delay in the processing of the second task due to dual-task interference and competition of two tasks for limited cognitive resources (Dell’Acqua et al., 2005; Sergent & Dehaene, 2004).

Overall, our results highlight that competition for attentional resources emerges as soon as the second task processing begins. The processing of the two tasks occurs in parallel during the decision-making process. Once the decision is made, there is another competition between the two tasks in motor areas for execution. This is evident from the reduced decoding accuracy, the temporal and conditional generalization patterns, and the DDM results (Figure 2-4). During this competition, intrinsic priorities and external instructions play a role in determining which task should be responded to first, allowing the winner to gain access to motor areas first. As a result, the reaction time of the second task is delayed compared to when there is no dual-task interference. In our study, we did not observe an above-chance decoding accuracy in the tone discrimination task. This is likely due to the subtle differences between the two tones, which do not allow us to detect the differences between the neural responses to the two tones in the subtle EEG signal. Future studies could use more distinct tones to examine changes in decoding accuracy for the tone task in a similar setting.

One interesting result obtained from the behavioral and MVPA was the significantly shorter reaction time and post-accumulation time, a smoother temporal generalization matrix, and quicker transfer of information to the motor cortex in the long SOA compared to the single lane-change condition. This indicates that once the tone task was presented, it worked as a cue for the lane-change task, triggering the deployment of attentional resources to the task. This result can be extended to actual driving situations in which sounding an alarm could draw attentional resources toward specific driving tasks, resulting in better performance and a decreased rate of accidents (Cheng et al., 2021).

The use of MVPA on the EEG data with excellent temporal resolution as well as the combination with the DDM, allowed us to successfully investigate the time course of the brain processes during dual-task interference. Consistent with hybrid theory, brain processes of two tasks occur in partial parallel to accumulate sensory evidence until a decision bound is reached. The relative saliency, the level of difference of conditions in each task, and the order of presentation affect the rate of decision accumulation. Once the decision is made, there is another competition in motor areas for execution. These results could inform future modeling efforts and can be used to improve driver assistant technologies to reduce the risk of accidents due to dual-task interference in real-life driving scenarios.

## Materials and methods

### Participants

The study involved human participants. Nineteen right-handed volunteers (9 females), 20-30 years old, with normal or corrected-to-normal vision and no history of neurological or psychiatric disorders, participated in the experiment. Additionally, all participants were not expert video game players, as defined by having less than 2 hours of video-game usage per week in the past year. Volunteers completed a consent form before participating in the experiment and were financially compensated after finishing the experiment. Experiments were approved by the ethics committee at the Institute for Research in Fundamental Sciences (IPM).

### Stimuli and Procedure

The dual-task paradigm consisted of tone discrimination and driving lane change tasks. Participants sat 50 cm from a 22″ LG monitor with a refresh rate of 60 Hz and a resolution of 1920 × 1080 and responded to the tasks using a computer keyboard. The driving environment was designed using the Unity 3D game engine.

For the tone task, a single pure tone of either a high (800 Hz) or low (400 Hz) frequency was presented for 200 milliseconds. Participants pressed the “x” and “z” keys on the computer keyboard with the middle and index fingers of their left hand to determine whether the tone was a high frequency or a low frequency, respectively. To provide feedback, if participants responded incorrectly, the green fixation cross turned red.

The driving environment included a straight highway with infinite lanes on the two sides, without left/right turns or any hills and valleys. This was to equalize all trials in terms of visual appearance for the experiment (Fig. 1A). The driving stimulus was composed of two rows of traffic cones with three cones in each row. In each trial, traffic cones were unexpectedly displayed on one side of the lanes, and the participants had to guide the vehicle to the lane with the cones and drive through them. The distance between the two rows of cones allowed the vehicle to drive through them without collisions easily. The cones were always presented in the lane immediately to the left or right of the driving lane, so the participants had to change only one lane per trial. The lane change was performed gradually, and the participants had to hold the corresponding key to direct the vehicle in between the two rows of cones and then release the key when the vehicle was in the correct situation. Any collision with the cones would be registered as an error, in which case the green fixation cross would disappear to provide negative feedback. The driving started with a given initial speed, which was kept constant. During the experiment, the participant moved to the right or left lanes by pressing the keys using the middle and index fingers of their right hand, respectively (Fig. 1A).

The experiment consisted of the dual-task and single-task conditions. In the dual-task trials, the two tasks were presented with either a short (100 ms) or long (600 ms) SOA. In the single-task trials, either the lane-change or the tone task was presented alone. In all dual-task trials, the tone task was presented first. There was a total of eight dual-task conditions: two SOAs (short and long) x two lane-change directions (shift right/shift left) x two tone conditions (low/high pitch); and four single-task conditions: two lane-change directions and two tone-discrimination conditions. In the dual-task conditions, the task presentation order was fixed so that the tone was always presented first, and the lane-change task was presented second. Each condition was repeated four times in each run, resulting in a total of 48 trials in every run. Each trial lasted 3s with an inter-trial interval varying from 1s to 3s. We used Optseq software (Dale et al., 1999) to optimize the presentation order of trials in each run. The participants completed 12 runs, each lasting 3.6 minutes (216 secs).

The participants were instructed to focus on the fixation cross at the center of the screen (Figure 1A) and first respond to tone and then the lane-change task as fast as possible. Using an Eyelink 1000 plus program, the eye tracking device was used to track their eye movements and remove trials exhibiting distracting eye movements.

The trial onset was determined by the presentation time of the tone stimulus, while the trial end was defined as the moment the car’s rear end reached the end of the traffic cone set. The lane-change task performance was evaluated as the percentage of trials in which the participant correctly detected the lane-change direction and successfully passed through the cones without any collisions. On the other hand, the tone task performance was assessed based on the number of correct identifications made by the participants.

Before the main experiment, all participants performed one training run similar to the main experiment. They would proceed to the main experimental runs if their performance were 80% or higher. All participants could reach this threshold.

### Drift-diffusion model

The drift diffusion Model (DDM) is a model that can be used to infer a wide range of cognitive decision processes (Gold & Shadlen, 2007; Ratcliff et al., 2016; Shadlen & Newsome, 2001). DDM assumes that participants accumulate information continuously until sufficient evidence is gathered to favor one of the choices (one of two thresholds is hit). The model uses the following partial differential equation (Shinn et al., 2020):

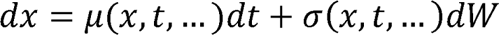

in which *µ*(*x*,*t*) is the instantaneous drift rate (i.e., the rate of evidence accumulation), σ (*sigma*) is the instantaneous noise, and *x*_0_ is the probability density function of the initial position (bias toward one of the decisions, here considered no bias for simplicity). The drift-diffusion process terminates when x reaches threshold A (Fig 1D).

Here, a variant of the drift-diffusion model (Zylberberg et al., 2012) was used, and another parameter *G (t,* μ *onset,* σ *onset)* was introduced. G is a cumulative Gaussian used to model the delayed onset of evidence accumulation and discount early samples compared to later samples. The reason for introducing this parameter was to observe if there was any delay in the accumulation of evidence due to divided attention at the beginning of the second task as the two tasks got closer in time.

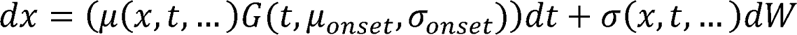

To solve DDM, Fokker-Planck equation, a partial differential equation using the probability density of the decision variable at position x and time t, was used as follows (Shinn et al., 2020; Sigman & Dehaene, 2005) :

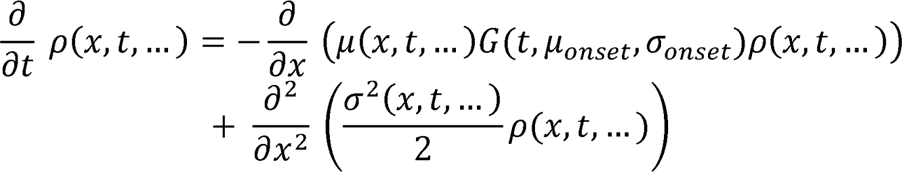

Subsequently the Fokker-Planck equation was approximated by discretization using the Crank-Nicolson method (Shinn et al., 2020) :

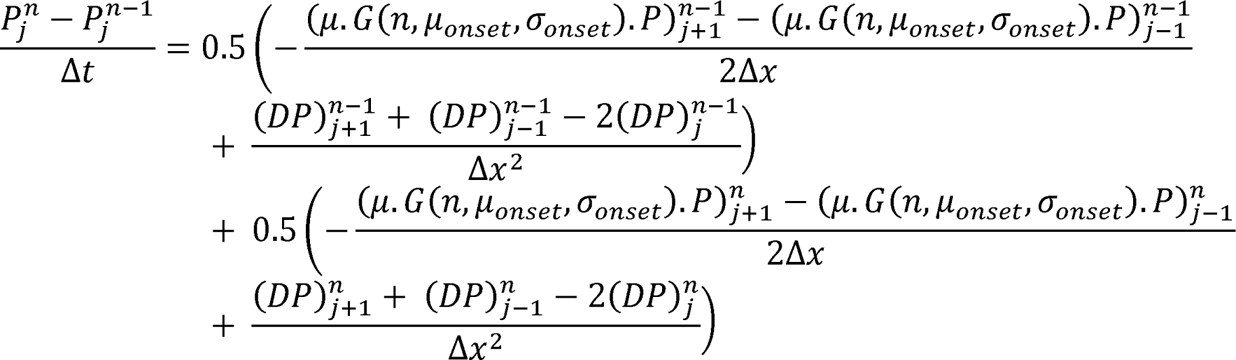

Here, 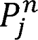 is defined as the probability distribution at the j^th^ space grid and the n^th^ time grid, and 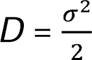 is the diffusion coefficient.

Taken together, four parameters were defined for each condition (short SOA, long SOA, and single lane change): two for the mean and standard deviation of *G* (cumulative Gaussian), one for drift rate, and another for post-accumulation time. The boundaries (*A*), sigma (σ), and starting point (*x*_0_) were considered equal among conditions.

The results of DDM can be highly affected by outliers (especially fast outliers). Due to positive skewness in the RT distribution, first, we log-transformed the distribution for each condition to remove fast outliers better. Then we used MATLAB (Mathworks, 2021a) function ‘’isoutlier’’ (which defines an outlier when a value is more than three scaled median absolute deviations (MAD) away from the median) to remove all outliers (Voss et al., 2015). Subsequently, data from all three conditions were pooled together, and the mentioned parameters were fitted using the maximum likelihood method. Bayesian adaptive direct search (BADS, (Acerbi & Ma, 2017)) in MATLAB was used as an optimization algorithm. In other words, the fitting procedure was done for all three conditions simultaneously with 14 parameters. Finally, R-squared was used to assess the model’s goodness of fit for each participant.

### EEG acquisition

The EEG signal was recorded using a 64-channel electrode cap with tin electrodes. The electrocap was organized according to the international 10/20 system and connected to a Bayamed reference amplifier. A mildly abrasive electrolyte paste reduced the impedance for all electrodes to less than 10kW. Data was sampled at 250Hz. The reference electrode was positioned on the left mastoid. The ground electrode was placed 1 cm inferior to the Oz electrode.

### EEG preprocessing

The EEG data was exported to EEGLAB 14b (Delorme & Makeig, 2004) in MATLAB for further preprocessing and analysis. Firstly, the data was re-referenced to an average reference and offline bandpass filtered between 1-40 Hz. The EEGLAB Cleanline method was then applied to eliminate line noise at 50 Hz and its subharmonics. To remove noisy channels, three automatic channel rejection methods of EEGLAB (spectrum, probability, and kurtosis with a z-score threshold of 5%) were utilized, and channels labeled as bad based on at least two methods were removed. Next, the raw data was epoched from -1 to 2.5 seconds relative to trial onset. A visual inspection was carried out to manually remove any non-stereotypical noise from the signals. The baseline was subtracted from the data using the remove epoch baseline method in EEGLAB. Following this, an ICA decomposition was performed using the ‘’runica’’ method (Delorme et al., 2007; Makeig et al., 2004). Components showing more than 90% probability for eye blink, muscle twitch, and channel noise were flagged for rejection. Additionally, components classified as residual gradient noise were manually removed. Epochs were again visually inspected to further eliminate residual noise before applying EEGLAB epoch rejection methods, which involved rejecting epochs with abnormal values exceeding 50 μV, improbable data with more than five standard deviations between channels, and improbable trends with a maximum slope/epoch of 50. Subsequently, the data was re-epoched from 500 ms prior to the onset of the tone to 1300 ms after the onset of the lane-change. For the single tone condition, data was epoched from 500 ms before the tone onset to 1500 ms after the tone onset. Eventually, the signal-to-noise (SNR) ratio was calculated for each participant based on epoched data, and participants for which SNR was lower than three were rejected. One subject was rejected based on SNR.

### Multivariate pattern analysis (MVPA)

To measure the temporal dynamics of dual-task information processing, we applied the MVPA analysis method to the EEG data for the lane-change and tone tasks, separately. In each task, three conditions of the short and the long SOAs and the single task were predefined. In the case of the tone discrimination task in each condition, MVPA was applied to data to assess whether information about tone discrimination (high vs. low) is encoded differently across conditions (Marti et al., 2015). Likewise, the same procedure was applied to the lane-change task to differentiate the left vs right shift in each condition. For each participant and each task, trials of the short and the long SOAs and single conditions were extracted from the EEG data. Then, data was normalized (Z-transformed baseline normalization) by their baseline of 200 to 0 ms prior to the stimulus onset.

Subsequently, for each task at each time point, a data matrix of the number of all trials x the number of sensors was created per condition. We first normalized the matrix (z-score of each channel-time feature) and then used a support vector machine (SVM) with a kernel function using MATLAB “fitcsvm” to decode right vs. left in the lane-change task and high vs. low in tone task, with a 10-fold stratified cross-validation approach. Stratified means the same proportion of each class was kept within each fold.

To compensate for fewer trials in single condition and get a reliable comparison, we applied a trial matching procedure. First, we randomly drew a set of trials from short and long SOAs with the same number of trials of a single condition with replacement. We proceeded by training and optimizing the classifier using a 10-fold cross-validation approach. This methodology was iterated 100 times, and the results were averaged across the 100 datasets to obtain the trial-matched decoding accuracy. Figure S2 also shows the results of long and short SOA decoding accuracy (which had comparable number of trials) without the resampling methods.

### Time-time decoding analysis

Time-time decoding accuracies were also obtained by cross-decoding across time. Each SVM classifier trained at a given time was tested at all other time-points and then averaged over the 10-fold cross validation, thereby demonstrating the cross-validated generalization across time. The complete ‘‘temporal generalization’’ (King & Dehaene, 2014) resulted in a matrix of training time * testing time.

### Generalization across Conditions

To evaluate how brain responses of each task were affected by the dual-task paradigm, the optimized SVM classifier for each condition at any given time was tested on its ability to successfully discriminate classes of each task at all the time samples of other conditions. The process was also executed through cross-validation and subsequently averaged to produce the mean classification accuracy across conditions. The outcome was training-time by testing-time matrix that was generalized across conditions.

### Searchlight analysis

To determine which electrode contributed most to the decoding performance in each time bin, searchlight analysis was used. In this analysis, time was the feature matrix, and the decoding performance of each electrode neighborhood was calculated. First, the square distance of electrodes was calculated using the X and Y positions of electrodes. Subsequently, electrodes for which this distance was less than 0.3 were considered neighbors and averaged together. Finally, a sliding window of 50 ms with steps of 4 ms was used, and at each step, the classification was carried out at the corresponding window. The MVPA-light package was used to compute decoding performance with nonlinear SVM as a classifier and a 10-fold stratified cross-validation method (Treder, 2020). A similar trial matching method was applied to the short and the long SOA conditions.

### Statistical Analyses

For each participant and each task, first the averaged cross-validated accuracy was calculated per condition at each time point. Then, we used the nonparametric Wilcoxon signed-rank test for random effect analysis. To determine time points with significant above-chance accuracy, we used a right-sided signed-rank test across participants (n = 18). To adjust p-values for multiple comparisons (e.g. across time), we further applied the false discovery rate (FDR) correction (FDR q < 0.05) (Groppe, 2024). In addition, to determine whether two conditions were significantly different at any time interval, we used a two-sided signed-rank test, FDR corrected across time.

Subsequently, nonparametric effect sizes are reported with an area under the curve (AUC) computed from the receiver operating curves (ROC) for searchlight analysis, temporal and conditional generalizations. AUC represents relative predictions of true positives (e.g., a trial was correctly classified as left/right in lane-change task or high/low in tone task) and predictions of false positives (e.g., a trial was incorrectly classified). An AUC of 0.5 corresponds to chance level as it means that true and false positives are equiprobable. Conversely, an AUC of 1 means a perfect class prediction. We used signed-rank tests with a threshold set at alpha = 0.05 to assess whether classifiers could predict the trials’ classes above chance level (50%). A correction for multiple comparisons was then applied with a false discovery rate (FDR q < 0.05).

One-way repeated measures ANOVA followed by a Tukey HSD multiple comparison test was used to compare differences between the conditions for behavioral performance, reaction times, and estimated parameters of the drift-diffusion model. Greenhouse-Geisser correction was performed whenever sphericity had been violated.

### Significance testing

We measured the onset latency, offset latency, and duration of each classification time course for each participant. Onset latency: At each training time, onset latency was defined as the earliest time where performance became significantly above chance (i.e., AUC > 0.5) for at least five consecutive time bins (20 milliseconds). Mean and standard deviation (SD) for onset latencies were calculated by resampling the data 100 times. In each iteration, 18 participants with replacements were chosen from the data, and the onset was calculated with the aforementioned definition. Offset latency: Likewise, offset latency for each training time was defined as the earliest time where performance became non-significant for at least five consecutive time bins (20 milliseconds). Mean and SD for offset latencies were also calculated by resampling. Finally, duration was the difference between offset and onset latencies.

