## Supplemental figures and tables for "The Temporal Profile of Dual-task Interference in the Human Brain"

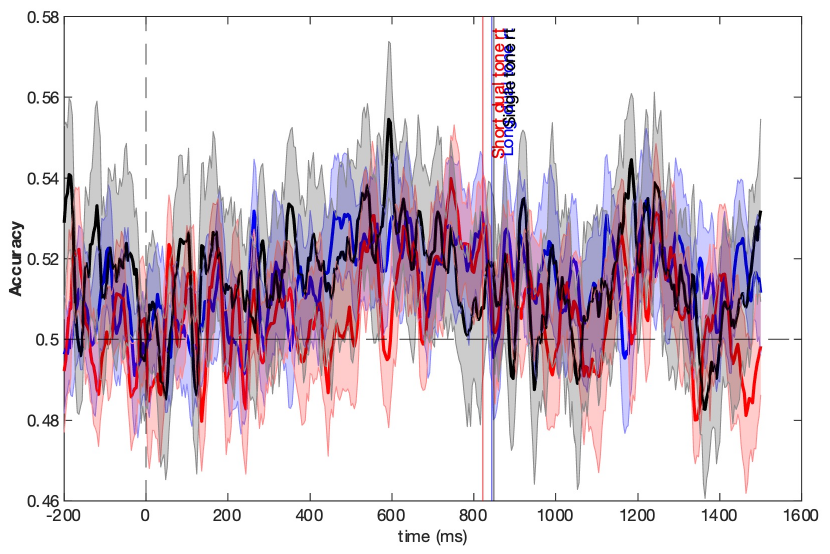

**Supplementary Figures**

**Figure S1. Temporal dynamics of dual-task interference in tone task.** For each subject, a nonlinear SVM as an MVPA method was trained in each time bin to discriminate between high vs low in tone task, and cross-validated accuracies were calculated and averaged across 18 subjects. As was mentioned, the decoding performance of the tone task was markedly compromised by the more salient driving task. As a result, there was no period of significantly above-chance performance in either of the conditions, nor their difference were significant. Shaded error bars represent SEM. The horizontal dashed line represents 50% accuracy corresponding to chance level. The vertical dashed line shows the stimulus (tone) onset. Three vertical lines are the corresponding reaction times of each condition

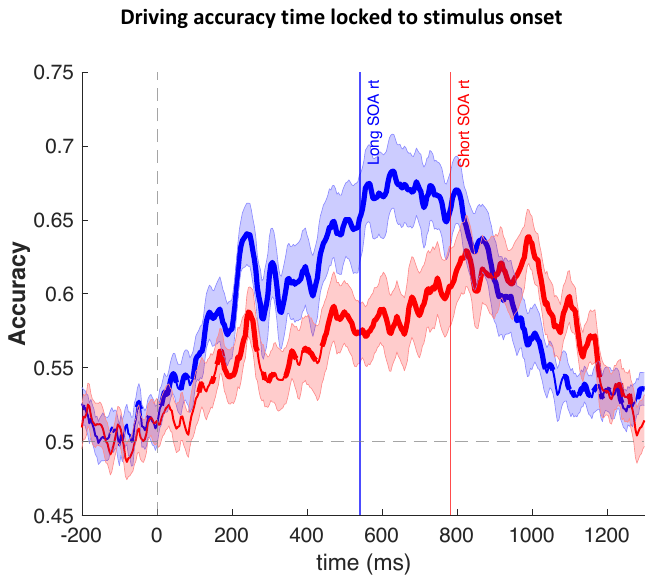

**Figure S2. Temporal dynamics of dual task interference in lane-change task without resampling method.** Multivariate pattern analysis of EEG data. Time courses of decoding accuracies for the long and the short SOA condition averaged across 18 subjects time-locked to the stimulus onset without the resampling method using all the trials for each subject. Dashed vertical lines show lane-change onset. The chance level decoding accuracy is shown by the horizontal dashed line at 0.5. Two vertical lines are the corresponding reaction times of long and short SOAs, respectively. Thicker lines indicate a decoding accuracy significantly above chance (right-sided signed-rank test, FDR corrected across time, q < 0.05), and shaded error bars represent the standard error of the mean (SEM). For display purposes, data was smoothed using a moving average with five sample points.

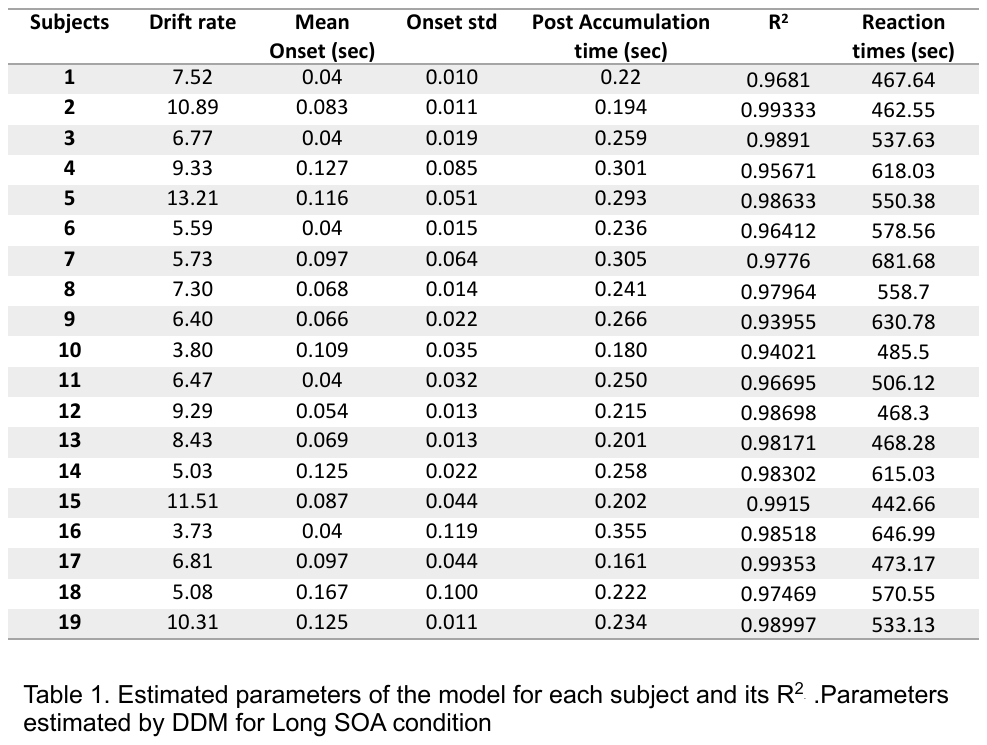

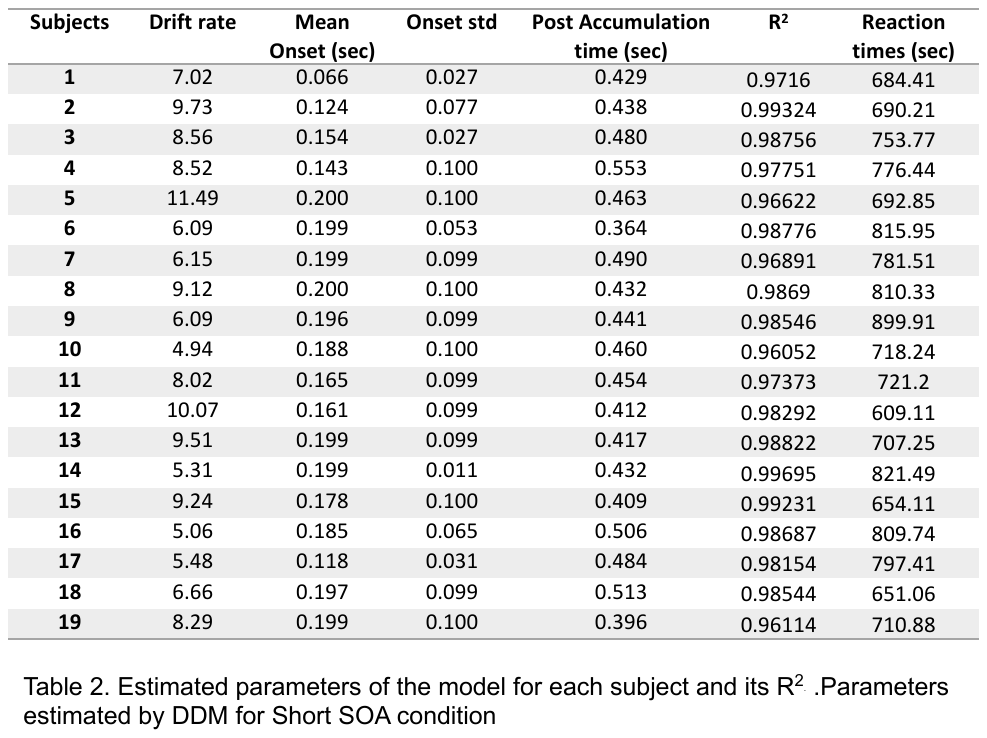

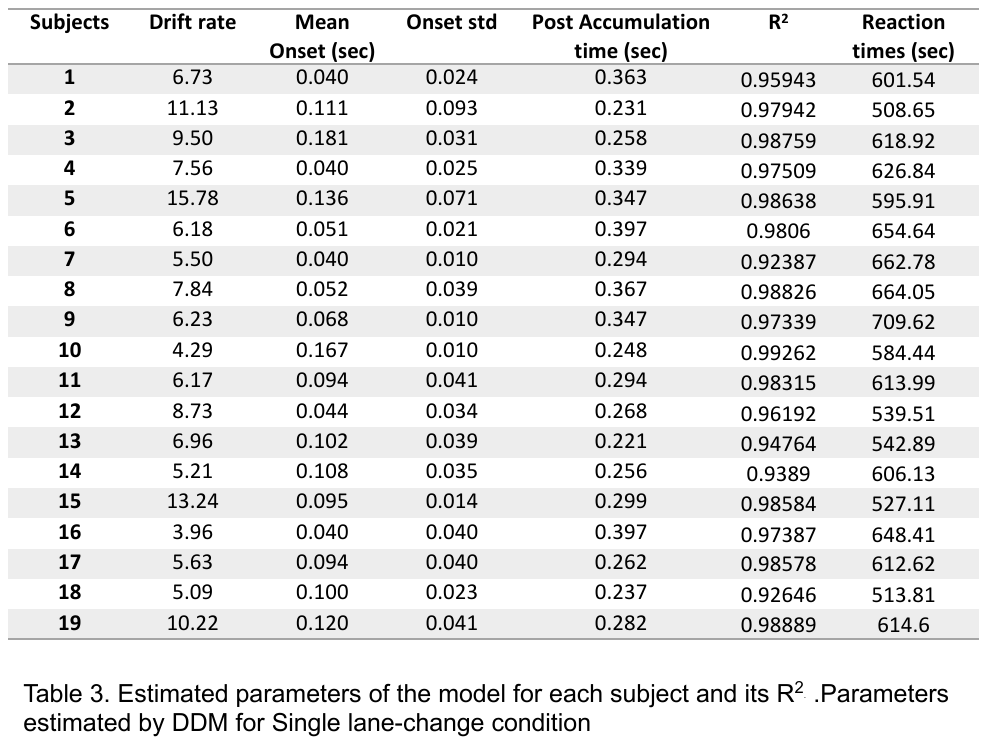
